## Supplemental Figures for "Completion of neural crest cell production and emigration is regulated by retinoic acid-dependent inhibition of BMP signaling"

Running title: Local RA signals the end of neural crest production

Key words: BAMBI, cell specification, dorsal interneurons, Foxd3, Hairy, neural tube, Raldh2, Roof plate, Snai2, Sox9.

**Abstract**

Production and emigration of neural crest cells is a transient process followed by the emergence of the definitive roof plate. The mechanisms regulating the end of neural crest ontogeny are poorly understood. Whereas early crest development is stimulated by mesoderm-derived retinoic acid, we report that the end of the neural crest period is regulated by retinoic acid synthesized in the dorsal neural tube. Inhibition of retinoic acid signaling in the neural tube prevents the normal upregulation of BMP inhibitors in the nascent roof plate and prolongs the period of BMP responsiveness which otherwise ceases close to roof plate establishment. Consequently, neural crest production and emigration are extended well into the roof plate stage. In turn, extending the activity of neural crest-specific genes inhibits the onset of retinoic acid synthesis in roof plate suggesting a mutual repressive interaction between neural crest and roof plate traits. Although several roof plate-specific genes are normally expressed in the absence of retinoic acid signaling, roof plate and crest markers are co-expressed in single cells and this domain also contains dorsal interneurons. Hence, the cellular and molecular architecture of the roof plate is compromised. Collectively, our results demonstrate that neural tube-derived retinoic acid, via inhibition of BMP signaling, is an essential factor responsible for the end of neural crest generation and the proper segregation of dorsal neural lineages.

Figure 1 – Figure Supplement 1

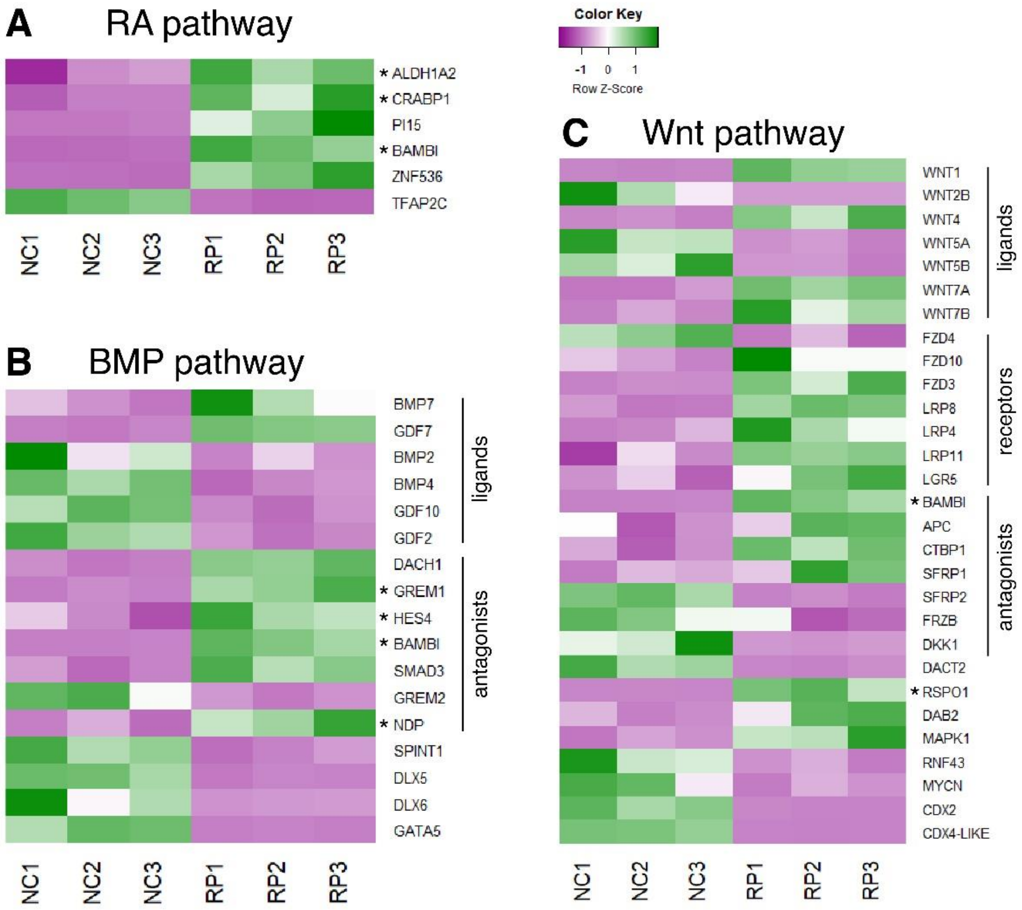


<u>Figure 1 – Figure Supplement 1 – Expression patterns of selected genes from a comparative RNA-seq analysis between NC and RP stages</u>

(A-C) Heat maps depicting the differential expression patterns of genes involved in RA (A), BMP (B) or Wnt (C) pathways, adapted from an RNA-seq analysis performed by Ofek et al. (Ofek et al., 2021) on premigratory NC (experimental triplicates NC1, NC2, NC3) and RP cells (experimental triplicates RP1, RP2, RP3). Genes discussed in the present study were marked by an asterisk.

### Figure 1 – Figure Supplement 2

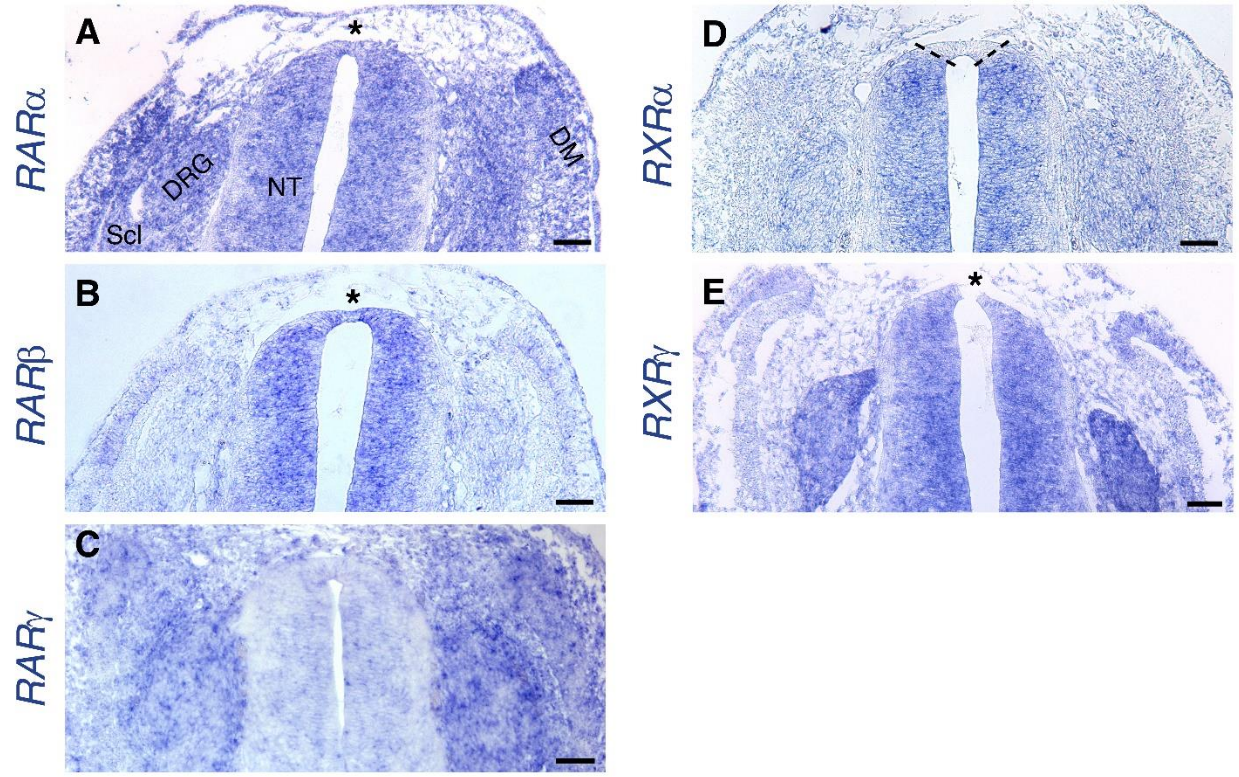


<u>Figure 1 – Figure Supplement 2 - Expression of RA receptors in the dorsal NT at the RP stage</u>

(A-E) ISH showing expression patterns of RA receptors at the RP stage (E4). [See also (Diez del Corral et al., 2003), for whole-embryo ISH at the NC stage]. Images were taken at somite levels 24-26.

(A) Expression of *RARα* is ubiquitous, including the dorsal NT (asterisk). (B) *RARβ* is expressed in the NT, including the dorsal domain (asterisk). (C) No significant expression of RARγ in the NT.

(D) *RXRα* is expressed throughout the NT, except for the dorsal domain (delineated by dashed lines). (E) Expression of *RXRγ* at the RP stage appears in the entire NT (asterisk dorsal to the RP) and in NC derivatives such as the DRG.

Abbreviations, DM, dermomyotome, DRG, dorsal root ganglion, NT, neural tube, No, notochord, Scl, sclerotome. Scale bar, 50 µm.

Figure 2 – Figure Supplement 1

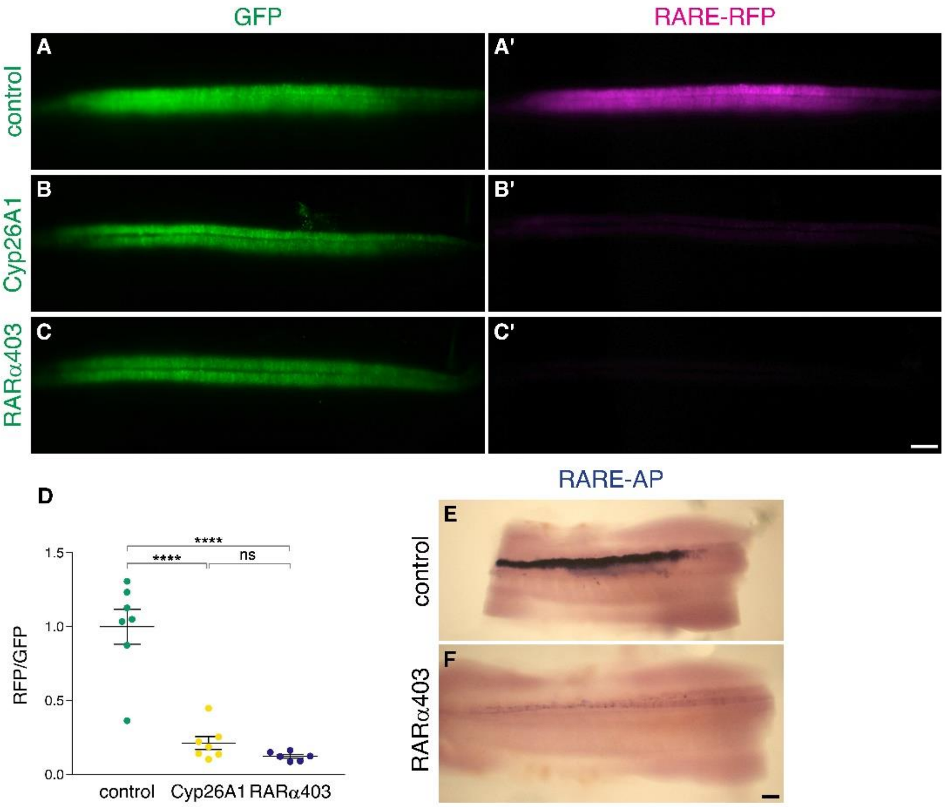


<u>Figure 2 – Figure Supplement 1 - Electroporation of Cyp26A1 or RARα403 into the NT effectively downregulates RA activity</u>

(A-D) Measurement of RA activity using RARE-RFP (magenta) in embryos electroporated with Cyp26a1, RARα403 or control GFP (green). NTs were co-electroporated bilaterally at E2 (23-24ss) and analyzed 30 hrs later. GFP was delivered along with the reporter construct and was used to assess the relative intensity of RFP fluorescence on whole-embryo images (D). N = 7, 7 and 6 embryos for control, Cyp26A1 and RARα403 groups, respectively. ****$p < 0.0001$, one-way ANOVA with post-hoc Tukey's tests.

(E-F) The same experimental conditions as in (A-D) were applied to unilaterally deliver RARE-AP (electroporated side facing the top). Note the robust AP staining in the control embryo, compared to the weak staining in the RARα403-treated embryo. N = 4/4 for both groups. Scale bar, 100 µm.

Figure 2 – Figure Supplement 2

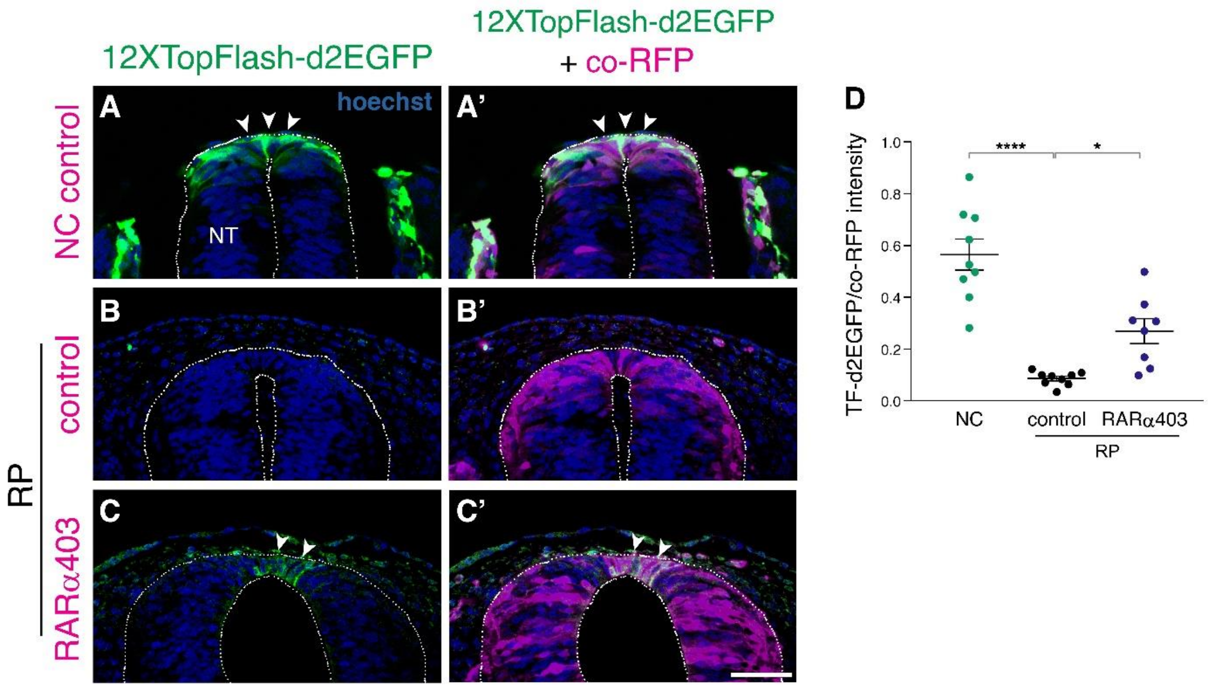


Figure 2 – Figure Supplement 2 - RA promotes the downregulation of Wnt signaling in the nascent RP

(A-C) A destabilized Wnt reporter, 12XTopFlash-d2EGFP, was electroporated bilaterally into the NT of E2.5 embryos (27 ss), and analyzed at E3 (35 ss, NC stage, control only) and at E4 (RP stage). RFP was co-electroporated to mark the electroporated region. Note the presence of Wnt activity in the premigratory NC (A-A', arrowheads) compared to its absence in the control RP (B-B'). Wnt activity is upregulated in RARα403-electroporated embryos compared to RFP only controls. Note the $GFP^+$ cells in the RARα403-treated embryo (C-C', arrowheads).

(D) Measurement of d2EGFP fluorescence signal intensity. N = 9 embryos for control NC and RP groups, and N = 8 for the RARα403-treated group, 12 sections were analyzed per embryo at somite levels 24-26. ****$p < 0.0001$, *$p < 0.05$, one-way ANOVA with post-hoc Tukey's test. Abbreviations, NT, neural tube. Scale bar, 50 μm.

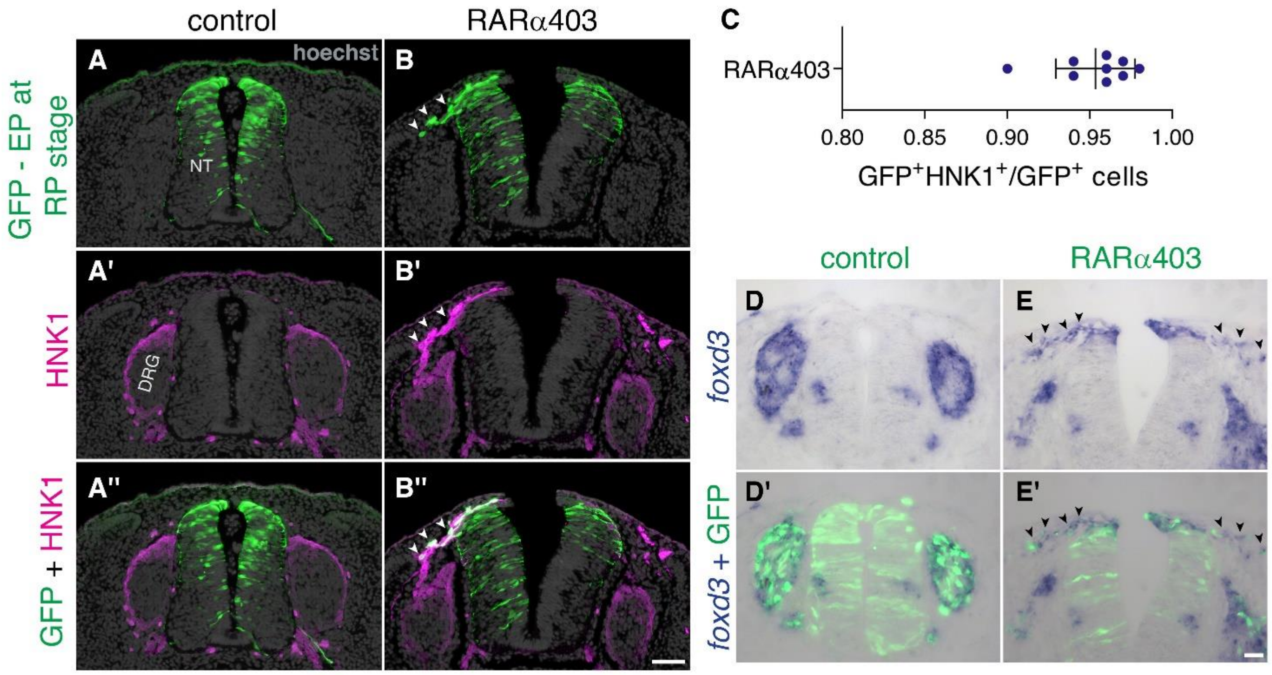


Figure 5 – Figure Supplement 1 - Late-delaminating cells display NC characteristics

(A-C) Immunostaining for HNK1 of embryos electroporated as in Fig. 5 (5, B-D) confirms that late-delaminating $GFP^{+}$ cells detected in RARα403-treated embryos exhibit traits specific to migrating NC cells (arrowheads in B-B''). The proportion of $GFP^{+}HNK1^{+}$ cells out of total $GFP^{+}$ cells outside the NT, is presented in (C). N = 9 embryos per treatment, 75 sections were monitored per embryo at somite levels 24-26. (D-E') ISH for *foxd3* of embryos electroporated as in Fig. 3 (3, A-B) reveals the presence of *foxd3*$^{+}$ NC cells in the mesenchyme adjacent to the dorsal NT of all RARα403-treated embryos (arrowheads in E-E'. N = 7). In contrast, only in 2/7 control embryos examined, few *foxd3*$^{+}$ cells were seen adjacent to the NT. In all control cases *foxd3*$^{+}$ cells are found in NC derivatives, such as the DRG (N=7). Abbreviations, DRG, dorsal root ganglion, NT, neural tube. Scale bar, 50 µm.

Figure 5 – Figure Supplement 2

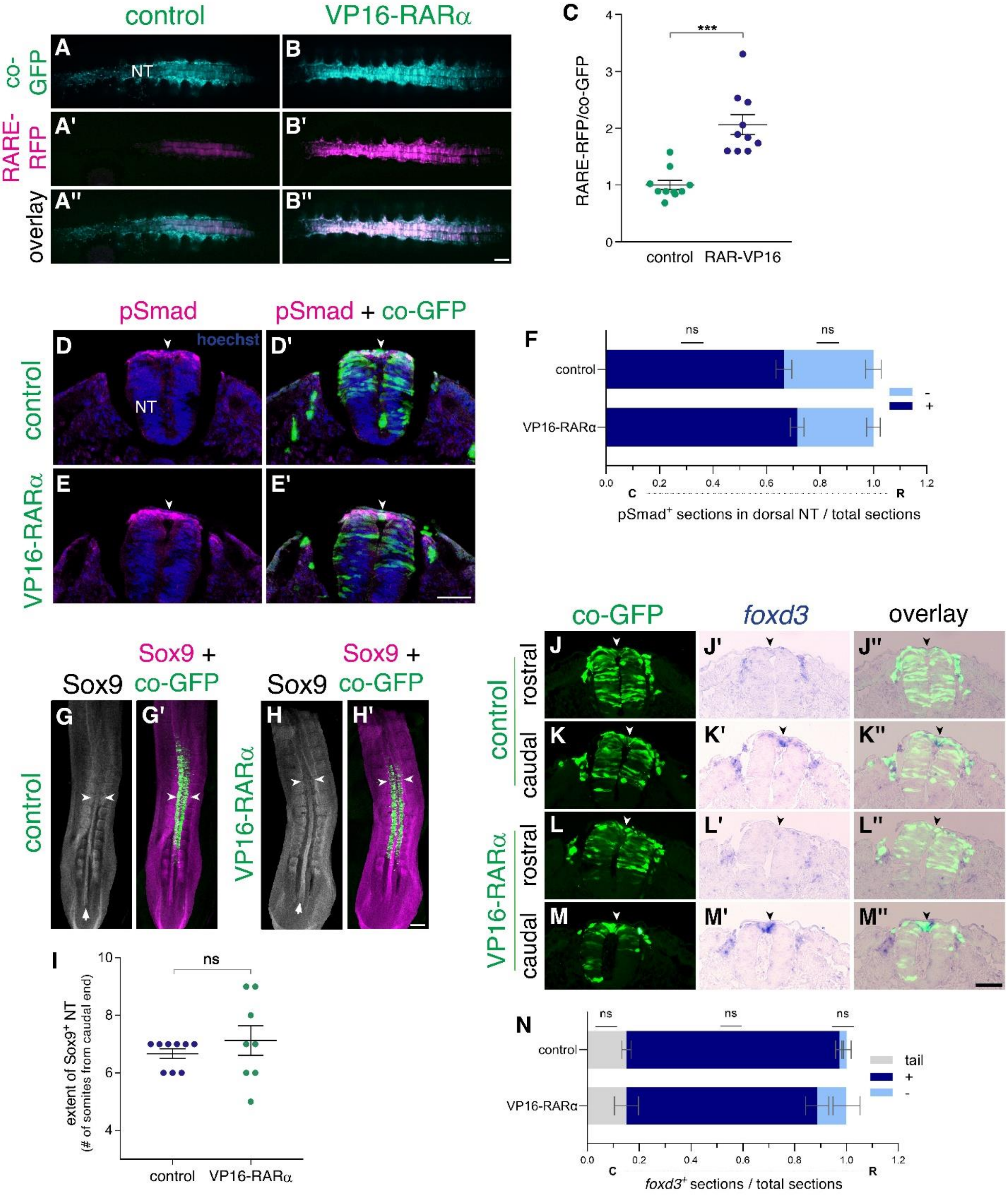


<u>Figure 5 – Figure Supplement 2 – Gain of RA function fails to prematurely downregulate BMP signaling and NC traits</u>

(A-C) VP16-RARα enhances RA signaling in the NT. Measurement of RA activity using RARE-RFP (magenta) in embryos electroporated with VP16-RARα or control plasmid (cyan). NTs were co-electroporated bilaterally at E2 (23-24ss) and analyzed 24 hrs later (E3). GFP was delivered along with the reporter construct and was used to assess the relative intensity of RFP fluorescence on whole-embryo images (C). N = 10 embryos for each group. ***p < 0.001, Welch's t-test.

(D-F) Dynamics of BMP activity assessed by immunostaining for pSmad1/5/9. Embryos were electroporated bilaterally at E2.5 (27ss) with either control PCAGG or VP16-RAR$\alpha$, together with GFP and analyzed 14 hrs later (35ss). (D-E') Images were taken from an axial level positioned just caudal to the level where downregulation of pSmad normally occurs in the dorsal NT midline of control embryos (somite level 23). Note that activation of RA signaling beyond normal levels failed to downregulate pSmad when compared to controls (arrowheads). (F) Quantification of the number of sections positive for pSmad (dark blue) serially monitored from tail to rostral end of the wing bud. Light blue represents sections with no detectable pSmad. Data are expressed as fraction of embryo length from tail. N = 6 and 8 embryos for control and VP16-RAR$\alpha$, respectively. Data were analyzed via Mann-Whitney test.

(G-I) The decreasing caudo-rostral gradient of Sox9 immunoreactivity in the NT is not affected by gain of RA function. Embryos were electroporated bilaterally at E2 (24ss) with either control PCAGG or VP16-RAR$\alpha$, together with GFP and analyzed at 28-29ss. Arrows in G,H depict expression of Sox9 in the NT, note also positive signal in adjacent somites. The axial level at which Sox9 staining in the dorsal NT is downregulated, was marked in whole-mount embryos (arrowheads) and quantified in (I) as the number of somites from the last-formed somite pair. N = 9 and 8 embryos for control and VP16-RAR$\alpha$, respectively, Mann-Whitney test.

(J-N) ISH for *foxd3* in embryos electroporated bilaterally at E2 (23-24ss) with either control PCAGG or VP16-RAR$\alpha$, together with GFP. Serial transverse sections were analyzed at 28-29ss in the caudal (C)-to-rostral (R) direction. (J-M") Images taken from axial levels positioned just rostral (J,L) and caudal (K,M) to *foxd3* downregulation (somite levels 23-24). In both treatments, *foxd3* was similarly expressed in the dorsal NT of caudal sections (K-K",M-M") and was absent more rostrally (J-J",L-L", arrowheads). (N) Quantification of serial sections expressing *foxd3.* Analysis in the caudo-rostral direction revealed a negative region in the tail (grey), followed by sections with positive signal (blue) and then exhibiting *foxd3* downregulation (light blue). Data are expressed as fraction of embryo length from the tail to caudal end of the wing bud. N = 9 and 6 embryos for control and VP16-RAR$\alpha$, respectively. Mann-Whitney test. Abbreviations, NT, neural tube. Scale bar, 50 $\mu$m (A-B",D-E',J-M"), 200 $\mu$m (G-H').

### Figure 5 – Figure Supplement 3

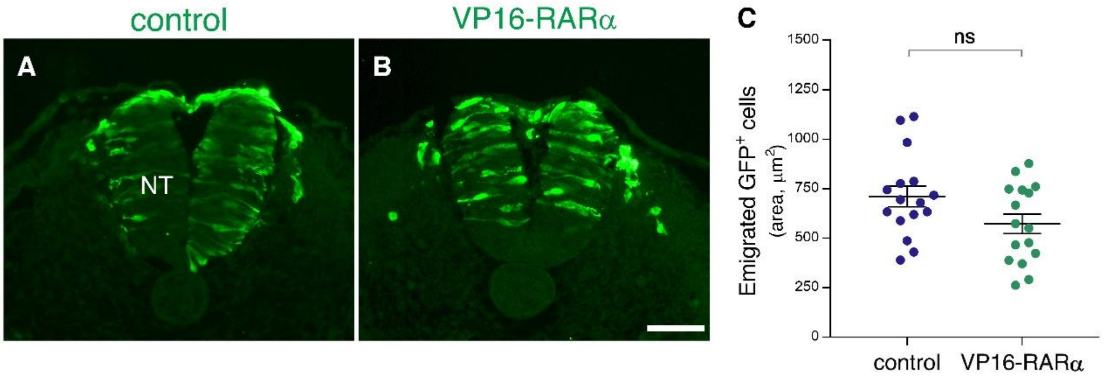


<u>Figure 5 – Figure Supplement 3 – No significant effect of VP16-RARα misexpression on completion of NC emigration</u>

(A-C) Emigration of transfected $GFP^+$ cells following gain of RA function. Embryos were electroporated bilaterally at E2.5 (27-28ss) with either control PCAGG or VP16-RARα, together with GFP and analyzed at somite levels 24-26 of embryos aged 35ss. Note ongoing emigration of $GFP^+$ cells in both treatments (A-B). (C) Quantification of NC emigration measured as the area containing $GFP^+$ cells outside the NT (21-42 sections per embryo). N = 16 embryos for each group. p=0.0654, Welch's t-test. Abbreviations, NT, neural tube. Scale bar, 50 μm.

Figure 6 – Figure Supplement 1

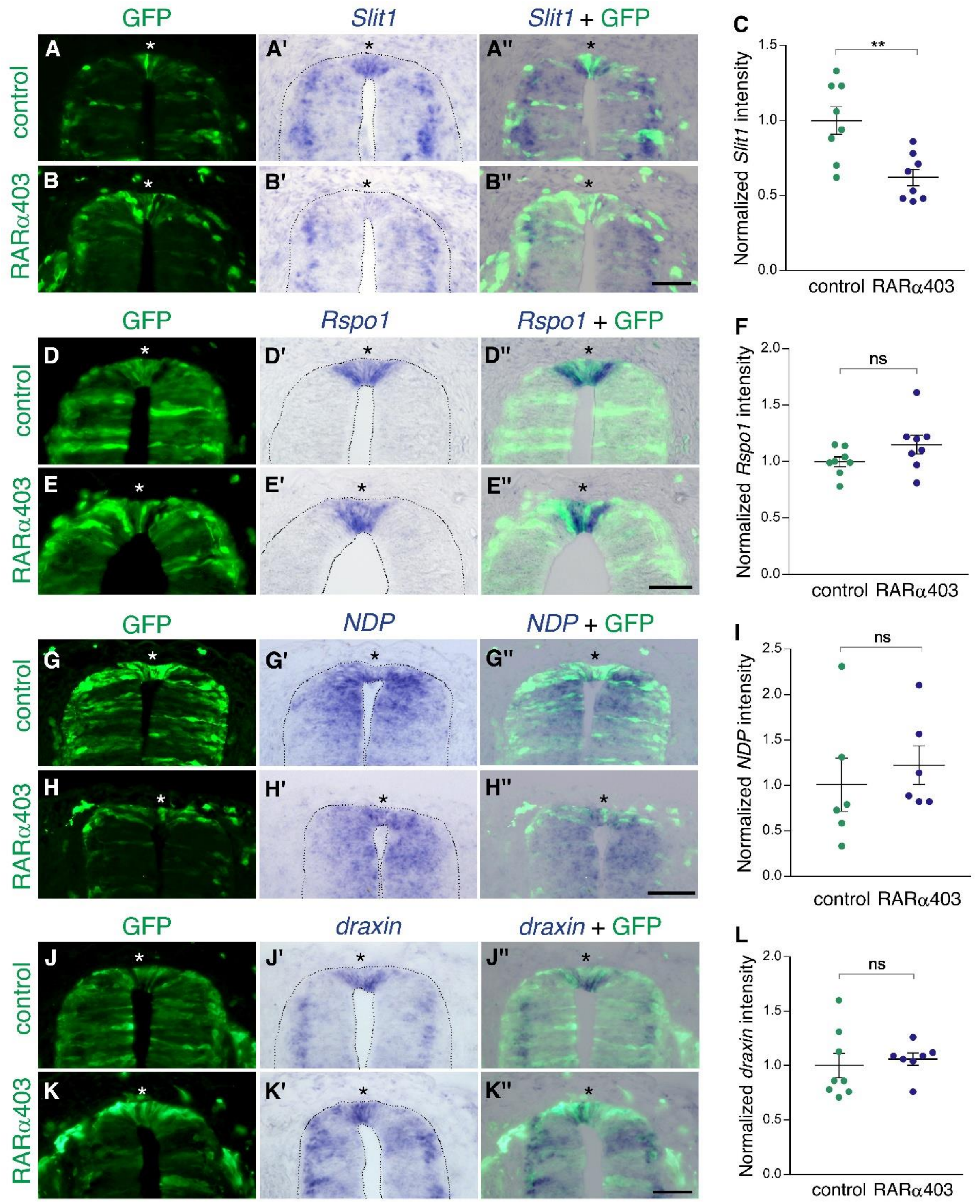


Figure 6 – Figure Supplement 1 - Inhibition of RA signaling only partially affects expression of definitive RP markers

ISH to monitor expression of RP-specific genes. Embryos were electroporated at E2.5 (27 ss) with either control PCAGG or RARα403, together with GFP, and analyzed at E4. Asterisks depict the RP domain. Imaging and analysis were performed at somite levels 25-27.

(A-C) ISH for *Slit1* reveals downregulation in the RP of RARα403-treated embryos. (C) Data quantification, N = 8 embryos for either control or RARα403 groups, 8-19 sections per embryo were analyzed. **$p < 0.005$.

(D-F) ISH for *Rspo1*. (F) Data quantification, N = 8 embryos for either control or RARα403 groups, 12-21 sections were analyzed per embryo.

(G-I) ISH for *NDP* (Norrin). (I) data quantification, N = 6 embryos for either control or RARα403 conditions, 6 sections were analyzed per embryo.

(J-L) ISH for *draxin*. (L) Data quantification, N = 8 control embryos and N =7 embryos that received RARα403; 10-17 sections were analyzed per embryo. Intensity comparisons of ISH staining presented in C, F, I and L were carried out using Welch's t-test. Scale bar, 50 µm.

Figure 6 – Figure Supplement 2

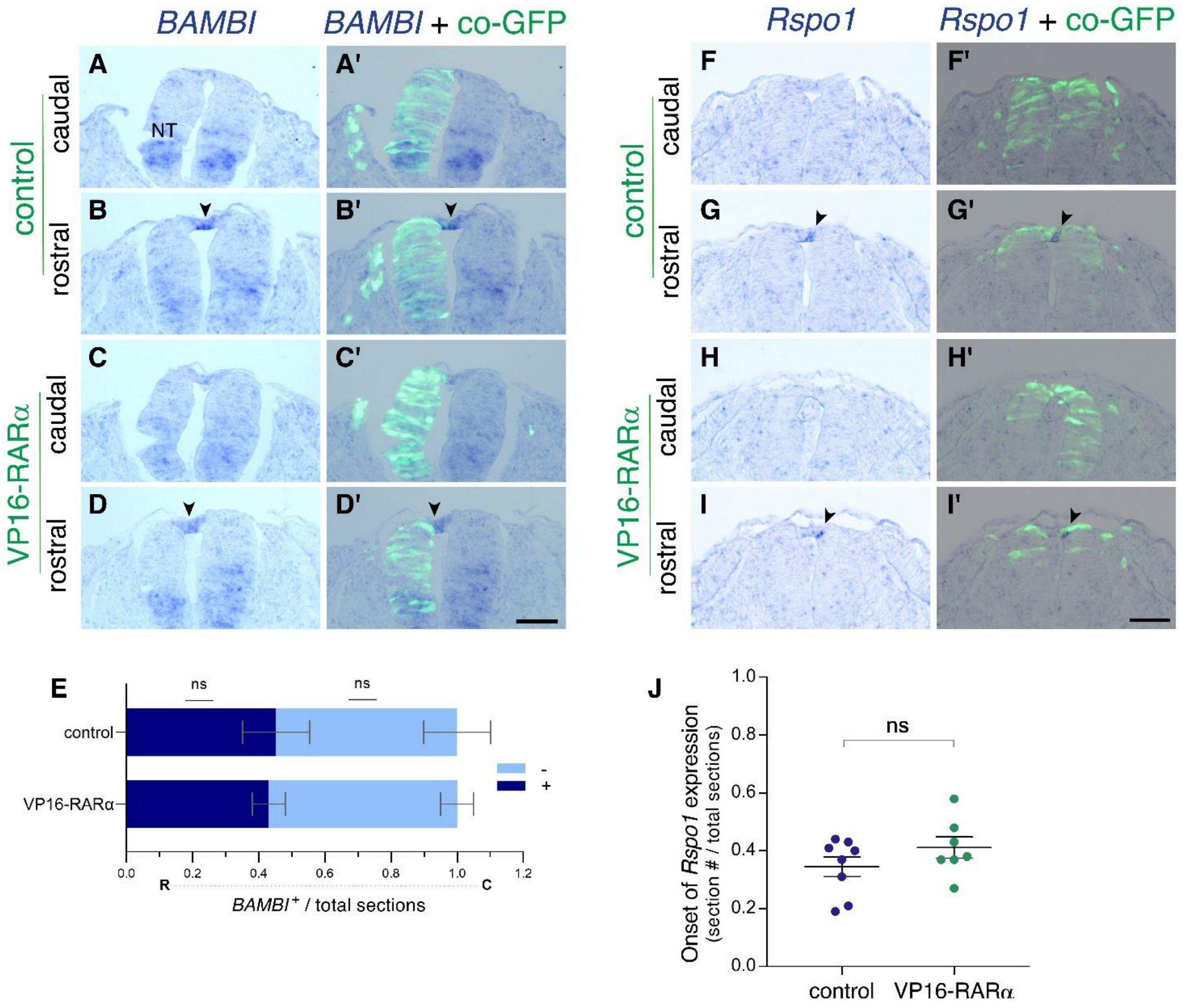


<u>Figure 6 – Figure Supplement 2 – Gain of RA signaling does not cause a premature upregulation of RP-specific genes</u>

(A-J) ISH for the RP-specific genes *BAMBI* and *Rspo1*, to assess their onset of expression in control and VP16-RARα–treated NTs.

(A-E) Embryos were electroporated unilaterally at E2 (22-24ss) with either control PCAGG or VP16-RARα, together with GFP and analyzed at 32-33ss for expression of *BAMBI*. (A-D') Images taken from axial levels positioned just rostral (B,D) or caudal (A,C) to the first appearance of *BAMBI* mRNA signal (somite level 25). Positive signal marked by arrowheads in B,D; no signal detected yet at levels of sections in A,C, highlighting similar expression dynamics of *BAMBI* in the dorsal NT in both treatments. (E) Quantification of serial sections expressing *BAMBI*. Analysis in the rostro (R)-caudal (C) direction revealed rostral sections with positive signal (blue) followed by caudal sections lacking *BAMBI* (light blue). Data are expressed as fraction of embryo length (# sections) from the caudal end of the wing bud to the tail. N = 6 and 5 embryos for control and VP16-RARα, respectively, Mann-Whitney test.

(F-J) Embryos were electroporated bilaterally at E2.5 (27-28ss) with either control PCAGG or VP16-RARα, together with GFP and analyzed at 35-36ss for expression of *Rspo1*. (F-I') Images taken from axial levels localized just rostral or caudal to the onset of *Rspo1* expression (somite level 28). Note that *Rspo1* begins to be faintly expressed in the rostral region in both treatments (G-G',I-I', arrowheads) but is absent from the caudal regions (F-F',H-H'). (J) Serial quantification of the first section with positive *Rspo1* signal out of the total number of sections. Data were monitored from the tail level to the rostral end of the wing bud. N = 8 and 7 embryos for control and VP16-RARα, respectively, Mann-Whitney test. Abbreviations, NT, neural tube. Scale bar, 50 μm.

Figure 7 – Figure Supplement 1

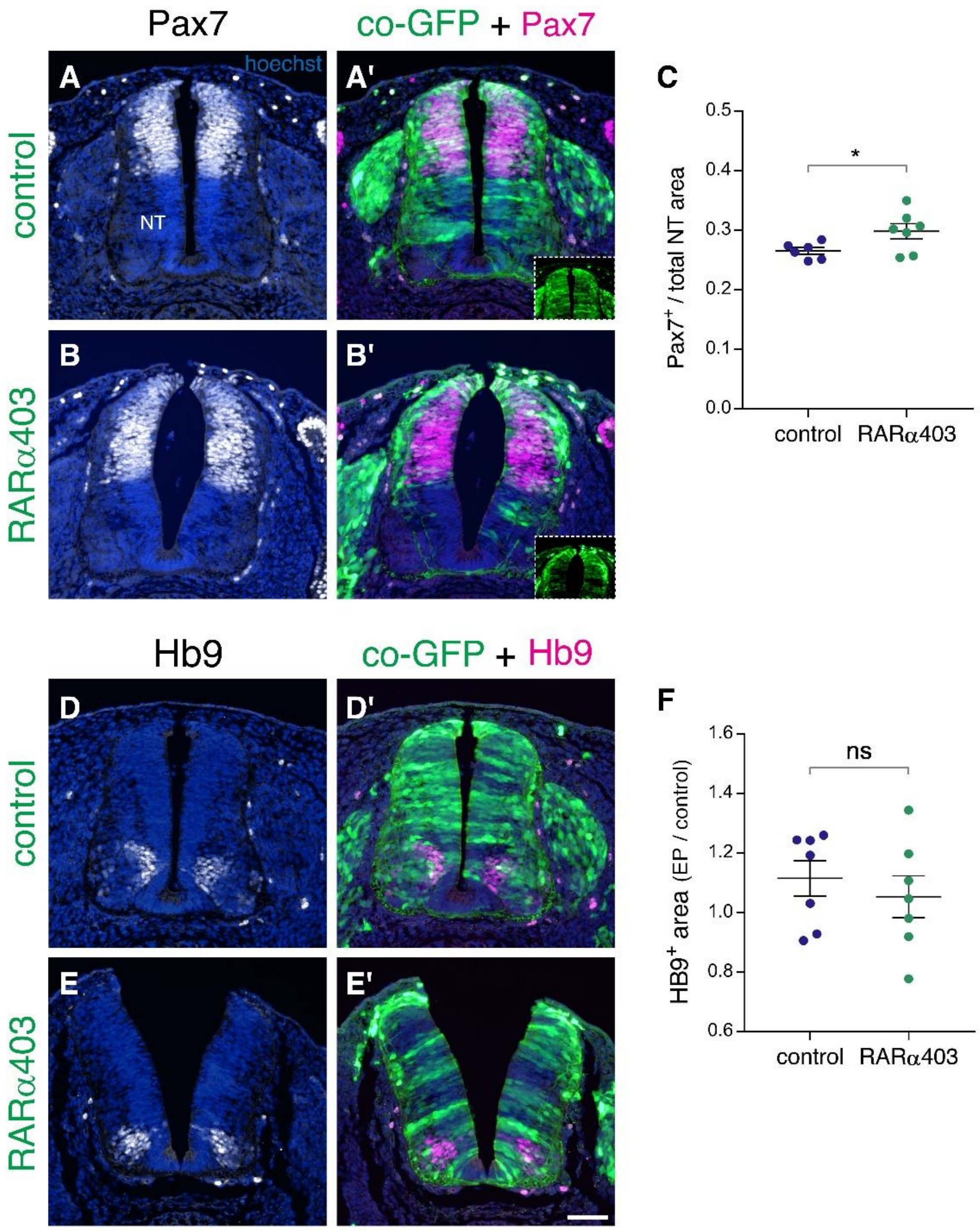


Figure 7- Figure Supplement 1- Loss of RA signaling during NC to RP transition does not affect dorso-ventral patterning of the NT

(A-F) Embryos were electroporated at E2.5 (27 ss) with control PCAGG or RARα403 along with GFP, and immunostained at E4 for Pax7 (A-C) or Hb9 (D-F).

(A-B') Staining for the dorsal marker Pax7. Insets in A' and B' show extent of electroporation. (C) Quantification of Pax7 staining. $Pax7^{+}$ area is expressed as a function of the total NT area. Ten sections per embryos were analyzed at the level immediately caudal to the upper limb. N = 6 and 7 for control and RARα403-treated embryos, respectively. *$p < 0.05$, Welch's t-test.

(D-E') Staining for the motoneuron marker Hb9. (F) Quantification of Hb9 staining. The area containing Hb9-positive cells was measured in 10 sections immediately caudal to the upper limb (electroporated region) and in 5 non-transfected sections at the upper limb level itself. Results are expressed as the ratio between electroporated /control areas of the same embryo. N = 7 for both control and RARα403-treated groups. $p = 0.52$ using Welch's t-test. Abbreviations, NT, neural tube. Scale bar, 50 μm.
